# Cognition’s dependence on functional network integrity with age is conditional on structural network integrity

**DOI:** 10.1101/2023.01.02.522493

**Authors:** Xulin Liu, Lorraine K Tyler, Cam-CAN, Simon W Davis, James B Rowe, Kamen A Tsvetanov

**Author notes:** Correspondence: Xulin Liu, University of Cambridge Department of Clinical Neurosciences, Herchel Smith Building, Cambridge Biomedical Campus, CB2 0SZ United Kingdom. Co-senior authors (James B Rowe, Kamen A Tsvetanov).

## Abstract

Maintaining good cognitive function is crucial for well-being across the lifespan. We proposed that the degree of cognitive maintenance is determined by the functional interactions within and between large-scale brain networks. Such connectivity can be represented by the white matter architecture of structural brain networks that shape intrinsic neuronal activity into integrated and distributed functional networks. We explored how the function-structure connectivity convergence, and the divergence of functional connectivity from structural connectivity, contribute to the maintenance of cognitive function across the adult lifespan. Multivariate analyses were used to investigate the relationship between function-structure connectivity convergence and divergence with multivariate cognitive profiles, respectively. Cognitive function was increasingly dependent on function-structure connectivity convergence as age increased. The dependency of cognitive function on connectivity was particularly strong for high-order cortical networks and subcortical networks. The results suggest that brain functional network integrity sustains cognitive functions in old age, as a function of the integrity of the brain’s structural connectivity.

## 1. Introduction

Maintaining good cognitive function is crucial for the preservation of well-being and quality of life across the lifespan (Beard et al., 2016; Sahakian, 2014). However, the degree of cognitive maintenance varies among individuals. The factors that maintain good cognitive function as we get older are still poorly understood. There is a pressing need to better understand the process of cognitive ageing in order to facilitate the maintenance of cognitive function in old age.

Brain function can be conceived in terms of interconnected networks, regionally specialized neural circuits that are integrated at multiple spatiotemporal scales (Breakspear & Stam, 2005). This network architecture can be characterized with connectomic approaches applied to structural or functional brain imaging data (Fornito, Zalesky, & Breakspear, 2015). Structural connectivity refers here to the white matter pathways between regions, which organize to enable energy-efficient processing (Betzel & Bassett, 2018; Park & Friston, 2013) and can be assessed by generating streamline estimates from diffusion-weighted MRI (Hermundstad et al., 2013). Structural connectivity typically declines with age (Betzel et al., 2014; Lim, Han, Uhlhaas, & Kaiser, 2015) and often strongly accounts for age-related changes in cognition (Davis, Szymanski, Boms, Fink, & Cabeza, 2019; Klein et al., 2016; Matejko, Price, Mazzocco, & Ansari, 2013; Moeller, Willmes, & Klein, 2015; Ponsoda et al., 2017). In contrast, functional connectivity refers to the node-to-node interactions between neurophysiologically active regions and can be estimated from functional magnetic resonance imaging (fMRI) using the temporal covariance between brain regional activity measured by blood oxygen level-dependent (BOLD) signal (Rosazza & Minati, 2011). Resting-state fMRI (rs-fMRI) can be used to characterize intrinsic and extrinsic connectivity of functional networks (Cole, Bassett, Power, Braver, & Petersen, 2014; S. M. Smith et al., 2009). Variation in resting-state functional connectivity is associated with ageing and cognitive performance (S. M. Smith et al., 2009). For example, as age increases, average functional connectivity within resting-state networks tends to weaken while functional connectivity between resting-state networks tends to increase (Betzel et al., 2014; L. Geerligs, Maurits, Renken, & Lorist, 2014; Guardia, Geerligs, Tsvetanov, Ye, & Campbell, 2022).

We have proposed that the maintenance of cognitive function with advancing age is increasingly dependent on the integrity of functional brain networks (Tsvetanov et al., 2016). This proposal replicates across cognitive states (Tomassini et al., 2022; Tsvetanov et al., 2018), neuroimaging modalities (Tibon et al., 2021), analytical approaches (Bethlehem et al., 2020; Linda Geerligs & Tsvetanov, 2017; Guardia et al., 2022) and in individuals with genetically increased risk of dementia (Chan et al., 2021; Passamonti et al., 2019; Rittman et al., 2019; Tsvetanov, Gazzina, et al., 2021). However, the increased reliance on functional integrity for maintaining cognitive functions in later life is poorly understood. For example, given that the interplay of structural and functional brain networks is important for cognition (Misic et al., 2016; Park & Friston, 2013; Vazquez-Rodriguez et al., 2019) and ageing (Davis, Kragel, Madden, & Cabeza, 2012; Davis et al., 2019; Persson et al., 2006), maintaining structural integrity may be critical for maintaining functional integrity in old age. Conversely, maintaining functional integrity may diverge from structural integrity, given the ability of the brain to reorganize despite brain structure loss (Cabeza et al., 2018), consistent with the unique mapping of functional connectivity on cognition (Zimmermann, Griffiths, & McIntosh, 2018). Modulating synaptic gains or nonlinear synchronous interactions, or both, may underpin this divergence (Park & Friston, 2013). Understanding this interaction is critical to anticipating the trajectory from healthy ageing to mild cognitive impairment and then to dementia, because the failure to preserve functional patterns in the face of inexorable declines in structural architecture is an example of poor brain maintenance (Cabeza et al., 2018). This trajectory is modulated by reserve and compensation which may both confer a degree of reorganization, but by different mechanisms (Barulli & Stern, 2013; Brickman et al., 2011; Driscoll & Troncoso, 2011). Reserve refers to a baseline capacity advantage, due to genetic and/or environmental factors such as education, enabling tolerance of later injury or degeneration; while compensation refers to the recruitment of additional neural resources that completely satisfy the increases in cognitive demands with age or disease progress even in the presence of significant structural atrophy (Cabeza et al., 2018). Separating functional connectivity that converges with structural connectivity from functional connectivity that diverges from structural connectivity would allow the exploration of brain maintenance mechanisms as well as evidence for reorganization.

Here we sought to better understand whether the increased reliance on maintaining functional integrity for good cognitive health in old age is (i) facilitated by structural network integrity, or (ii) independent of structural network integrity. A challenge is that neither structural and functional substrates of cognition nor the effects of ageing are mediated by a single connection. We therefore require multivariate integrative approaches to identify (i) shared signals between structural and functional connectomes (*function-structure connectivity convergence*) and (ii) unique signals in functional connectomes that are independent of differences in structural connectomes (*function-structure connectivity divergence*). We then tested whether signals associated with *function-structure connectivity convergence* and/or *function-structure connectivity divergence* became increasingly related to cognitive function with age.

## 2. Material and Methods

### 2.1 Cohorts and participants

The Cambridge Centre for Ageing and Neuroscience (Cam-CAN) cohort study recruited healthy adults, who were drawn from the general population via Primary Care Trust (PCT)’s lists within the Cambridge City (UK) area, in three stages (Shafto et al., 2014). In Stage 1, 3000 adults aged 18 and above were recruited for a home interview. In Stage 2, a subset of 700 participants aged 18-88 (100 per age decile) was selected to participate in neuroimaging (e.g., structural MRI and fMRI) and cognitive tests (Shafto et al., 2014; Taylor et al., 2017). We used data from Stage 2 participants in this study. Ethical approval was obtained from the Cambridge 2 Research Ethics Committee, and written informed consent was given by all participants. Participants performed a battery of cognitive tasks outside the scanner. A full description of cognitive tasks performed in the Cam-CAN cohort is described elsewhere (Shafto et al., 2014). Cognitive tasks examined in this study are described in the next section. We included 473 subjects in the analysis of structure-function connectivity correlation and the analysis of relationship between connectivity and age. The demographic characteristics of participants are reported in **Table 1**. Demographic variables were compared between age groups using one-way ANOVA for continuous variables and using the chi-square test for categorical variable.

**Table 1.** Characteristics of participants.

| | Age range | | | | | | | Difference<br>between deciles<br>(ANOVA or $\chi^2$) |
| --- | --- | --- | --- | --- | --- | --- | --- | --- |
|  | All | 18-27 | 28-37 | 38-47 | 48-57 | 58-67 | 68-81 |  |
| *n* | 473 | 38 | 97 | 93 | 88 | 88 | 69 |  |
| Mean age (years) | 48.9 | 23.4 | 32.9 | 42.8 | 52.3 | 62.4 | 72.2 |  |
| Gender |  |  |  |  |  |  |  |  |
| *n* (%) |  |  |  |  |  |  |  |  |
| Males | 243 | 22 (58) | 49 (51) | 48 (52) | 43 (49) | 46 (52) | 35 (51) | 0.94 |
| Females | 230 | 16 (42) | 48 (49) | 45 (48) | 45 (51) | 42 (48) | 34 (49) |  |
| Addenbrooke's<br>Cognitive Examination-Revised (ACE-R) |  |  |  |  |  |  |  |  |
| Mean ± SD | 95.7±4.0 | 95.4±5.1 | 97.2±3.2 | 96.4±3.3 | 96.0±3.6 | 95.6±4.1 | 92.8±4.5 | < 0.0001 |
| Cattell score |  |  |  |  |  |  |  |  |
| Mean ± SD | 33.3±5.9 | 37.7±3.9 | 37.1±4.3 | 35.1±4.3 | 33.4±4.4 | 30.6±5.4 | 26.8±6.0 | < 0.0001 |

### 2.2 Cognitive assessment

The cognitive tests of interest in this study spanned major cognitive domains and intelligence, including the Cattell culture fair test as a measure of fluid intelligence (Cattell, Cattell, Institute for, & Ability, 1960), the spot-the word test (Baddeley, Emslie, & Nimmo-Smith, 1993) as a measure of crystallized intelligence, visual short-term memory as a measure of working memory, motor response consistency (i.e., the inverse of response variability) as a measure of response consistency, the Hotel task as a measure of multitasking (Manly, Hawkins, Evans, Woldt, & Robertson, 2002), and Benton faces as a measure of face recognition (Benton, 1983). These cognitive domains have been shown to be increasingly associated with functional connectivity in older adults (Tsvetanov et al., 2016) and therefore were analyzed as the main cognitive tasks of interest in this study. Subjects with missing cognitive data (*n* = 48, female/male = 30/18; age range 19-77, age mean±SD = 48.4±17.6) were excluded in the analysis of relationship between connectivity and cognition.

### 2.3 Image acquisition and processing

#### 2.3.1 T1 structural MRI

Imaging data from Cam-CAN were acquired using a 3T Siemens TIM Trio. A 3D structural MRI was acquired using T1-weighted sequence with generalized autocalibrating partially parallel acquisition with acceleration factor 2; repetition time (TR) = 2250 ms; echo time (TE) = 2.99 ms; inversion time (TI) = 900 ms; flip angle α = 9°; field-of-view (FOV) = 256 × 240 × 192 mm; resolution = 1 mm isotropic; acquisition time of 4 min and 32 s.

Preprocessing of T1-weighted images used standardized preprocessing consistent with Cam-CAN data processing protocol (Taylor et al., 2017; Tsvetanov, Henson, et al., 2021). The Automatic Analysis (Cusack et al., 2014) pipelines implemented in Matlab (MathWorks) were used. The T1 image was initially coregistered to the MNI template, and the T2 image was then coregistered to the T1 image using a rigid-body transformation. The coregistered T1 and T2 images were used in a multichannel segmentation to extract probabilistic maps of six tissue classes: grey matter, white matter, cerebrospinal fluid, bone, soft tissue, and residual noise. The native space grey matter and white matter images were submitted to diffeomorphic registration (DARTEL) (Ashburner, 2007) to create group template images. Each template was normalized to the MNI template using a 12-parameter affine transformation. Images were modulated to correct for individual brain size.

#### 2.3.2 Diffusion-weighted imaging (DWI)

DWI data were processed using FSL (https://fsl.fmrib.ox.ac.uk/fsl/fslwiki; RRID:SCR_002823) and Mrtrix (http://mrtrix.org). Data were denoised, corrected for eddy currents, and bias-field corrected. A constrained spherical deconvolution (CSD) model was used in calculating the fiber orientation distribution, which was used along with the brain mask to generate whole-brain tractography (seed = at random within mask; step size = 0.2 mm; 10 million tracts). After tracts were generated, they were filtered using spherical-deconvolution informed filtering of tractograms (SIFT) to improve the quantitative nature of the whole-brain streamline reconstructions (R. E. Smith, Tournier, Calamante, & Connelly, 2013). This algorithm improves the selectivity of structural connections by determining whether a streamline should be removed using a cost function estimated on the difference between a) the number of fibers obtained from the fiber orientation distribution, and b) the proportionate amplitude of the fiber orientation distributions. Tracts were processed by SIFT until 1 million tracts remained.

#### 2.3.3 Resting-state fMRI

For rs-fMRI, echoplanar imaging (EPI) acquired 261 volumes with 32 slices (sequential descending order, slice thickness of 3.7 mm with a slice gap of 20% for whole-brain coverage, TR = 1970 ms; TE = 30 ms; flip angle α = 78°; FOV = 192 mm × 192 mm; resolution = 3 mm × 3 mm × 4.44 mm) during 8 min and 40 s. Participants were instructed to lie still with their eyes closed. The initial six volumes were discarded to allow for T1 equilibration. We quantified participant head motion using the root mean square volume-to-volume displacement (Jenkinson, Bannister, Brady, & Smith, 2002). The imaging data were preprocessed using Automatic Analysis (Cusack et al., 2014) calling functions from SPM12. EPI data preprocessing included (1) spatial realignment to correct for head movement and movement by distortion interactions, (2) temporal realignment of all slices to the middle slice, and (3) coregistration of the EPI to the participant’s T1 anatomical scan. The normalization parameters from the T1 stream were applied to warp functional images into MNI space.

Resting-state fMRI data were further processed using whole-brain independent component analysis (ICA) of single-subject time series denoising, with noise components selected and removed automatically using the ICA-based Automatic Removal of Motion Artifacts toolbox (AROMA) (Pruim, Mennes, Buitelaar, & Beckmann, 2015; Pruim, Mennes, van Rooij, et al., 2015). This was complemented with a general linear model detrending of the fMRI signal (L. Geerligs, Tsvetanov, Cam, & Henson, 2017), covarying out six realignment parameters, white matter and cerebrospinal fluid signals, their first derivatives and quadratic terms. Global white matter and cerebrospinal fluid signals were estimated for each volume from the mean value of white matter and cerebrospinal fluid masks derived by thresholding SPM tissue probability maps at 0.75. Data were band-pass filtered (0.0078–0.1 Hz) using a discrete cosine transform (Bright, Tench, & Murphy, 2017; Hallquist, Hwang, & Luna, 2013; Lindquist, Geuter, Wager, & Caffo, 2019).

### 2.4 Structural connectivity

Structural connectomes were generated by using FLIRT to apply a linear registration to the Brainnetome atlas (Fan et al., 2016) to register it to each subject’s native diffusion space. The Brainnetome atlas was developed to link functional and structural characteristics of the human brain (Fan et al., 2016) and provides a fine-grained whole-brain parcellation with a superior representation of age-related differences in brain structure compared to other cortical parcellation schemes (Long et al., 2018; Madan & Kensinger, 2018). Structural connectivity was defined as the number of streamlines connecting any pair of regions obtained from the parcellation schemes. The vectorized upper triangle of the connectivity matrices from 473 subjects was used.

Structural connectivity is susceptible to the occurrence of false positive and false negative connections and therefore a group threshold is often applied (de Reus & van den Heuvel, 2013). We applied a threshold of 50% which is within the optimal range recommended to eliminate false positive and false negative connections (de Reus & van den Heuvel, 2013). Structural connections that were 0 for 50% of subjects or more were not included in the analysis, resulting in 7934 remaining connections for SC. To ensure that thresholding did not affect the variables of interest significantly, we performed sensitivity analysis investigating the correlation of unthresholded structural connectivity (i.e., 30135 connections) with age and with functional connectivity.

### 2.5 Functional connectivity

Functional connectivity was estimated based on Pearson’s correlation of post-processed rs-fMRI time series between each pair of the 246 brain nodes, where the nodes were defined using the Brainnetome atlas (Fan et al., 2016). A vectorized connectivity matrix (upper triangle) was created for each subject containing 30135 connections. Functional connectivity corresponding to the structural connectivity after thresholding was included, resulting in 7934 connections for functional connectivity. We refer to this functional connectivity as *functional connectivity observed*. To ensure that thresholding did not affect the variables of interest significantly, we also investigated the correlation of unthresholded *functional connectivity observed* (i.e., 30135 connections) with age and with structural connectivity.

### 2.6 Statistical analysis

#### 2.6.1 Function-structure connectivity convergence and function-structure connectivity divergence

Canonical correlation analysis (CCA) (Correa, Li, Adali, & Calhoun, 2008; Hotelling, 1936) with 10-fold cross-validation was used to investigate the association between *functional connectivity observed* and structural connectivity. CCA is a data-driven multivariate analysis method that maximizes the correlation between two datasets while minimizing redundancy across variables within datasets (Hotelling, 1936). Structural connectivity was log-transformed and both *functional connectivity observed* and structural connectivity were normalized. A structural connectivity matrix (connections by subjects, 7934 × 473) and a functional connectivity observed matrix (connections by subjects, 7934 × 473) were entered into CCA to identify spatial components that were significantly correlated between these two matrices. All significant components were regressed out from the original *functional connectivity observed* matrix to create a remaining matrix (i.e., to determine the residuals in the *functional connectivity observed* after regressing out the signals identified by CCA). We referred to the residual signals as *function-structure connectivity divergence* in this paper (connections by subjects, 7934 × 473). The functional connectivity that was significantly correlated with structural connectivity was obtained by correlating the original *functional connectivity observed* with the significant function-structure connectivity correlated components resulted from CCA mentioned above, and this functional connectivity is referred to as *function-structure connectivity convergence* (connections by subjects, 7934 × 473) in this paper. The complete analytical processes are illustrated in **Figure 1**.

**Figure 1.**
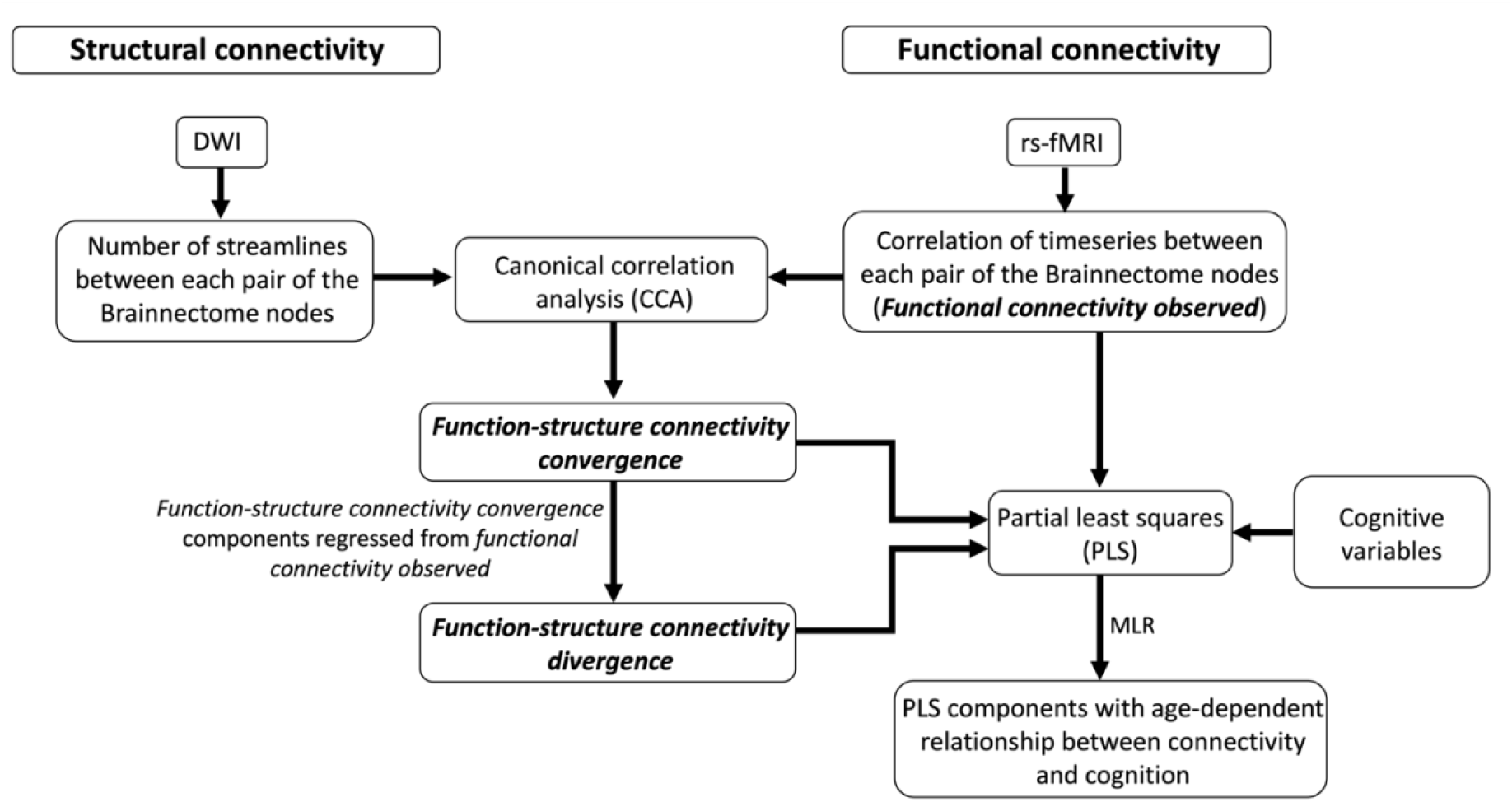
Summary of the analytical process of brain connectivity in relation to cognition. Abbreviations: DWI, diffusion-weighted imaging; rs-fMRI, resting-state functional magnetic resonance imaging; MLR, multiple linear regression.

#### 2.6.2 Relating connectivity to age

Connectivity was mapped onto 7 cortical networks (visual, somatomotor, dorsal attention, ventral attention, limbic, frontoparietal and default) as defined by Yeo et al (Yeo et al., 2011), as well as a subcortical set of regions which includes the amygdala, hippocampus, basal ganglia and thalamus. The resulting connectivity matrix contained 8 within-network connections and 28 between-network connections.

Multiple linear regression with robust fitting algorithm (Matlab function fitlm.m) was used to investigate the correlation of *function-structure connectivity convergence* and *function-structure connectivity divergence* with age for each of the 36 network connections. Gender and head motion were included as covariates of no interest. The model’s formula took the following form using Wilkinson notation (Wilkinson & Rogers, 1973): Age ∼ *Function-structure connectivity convergence/divergence* + Gender + Head motion. For comparison, the correlation of *functional connectivity observed* with age was also investigated using the multiple linear regression model: Age ∼ *Functional connectivity observed* + Gender + Head motion. All variables were normalized before entering into the regression model. A false discovery rate (FDR) corrected *P*-value < 0.05 was considered statistically significant.

#### 2.6.3 Relating connectivity to cognition

We included subjects with data recorded for all 6 cognitive variables of interest and therefore 425 subjects were included in the cognition analysis. We performed a multivariate analysis using partial least squares (PLS) which optimizes the covariance between datasets (McIntosh, Bookstein, Haxby, & Grady, 1996). Although both CCA and PLS are useful to characterize relationships between two datasets, PLS has been suggested as a more suitable method for mixed datasets (Beaton, Saporta, & Abdi, 2020; Grellmann et al., 2015), especially in the case of brain-cognition relationships (Mihalik et al., 2022). PLS with 10-fold cross-validation was used to investigate the correlation (i) between *functional connectivity observed* and cognitive variables, (ii) between *function-structure connectivity convergence* and cognitive variables, and (iii) between *function-structure connectivity divergence* and cognitive variables. Significant components (*P* < 0.05) were identified from PLS. To better understand whether PLS’s function-cognition relationships varied with age, the connectivity subject scores and cognition subject scores of the significant components were modelled in a moderation analysis with age using multiple linear regression. Variables were analysed in the following model using Wilkinson’s notation: Cognition ∼ Network connectivity*Age + Gender + Head motion. All variables were normalized before entering into the regression analysis. An FDR-corrected *P*-value < 0.05 was considered statistically significant.

## 3. Results

### 3.1 Functional connectivity observed, Function-structure connectivity convergence and function-structure connectivity divergence

The structural connectivity and functional connectivity (average connectivity across all regions for each subject) decreased with age (**Figure 2**). The correlation between structural connectivity and functional connectivity also decreased with age (**Figure 2**). The results for unthresholded data were consistent with the results for thresholded data, suggesting that thresholding of the connectivity data does not introduce major bias in our findings (**Figure 2**).

**Figure 2.**
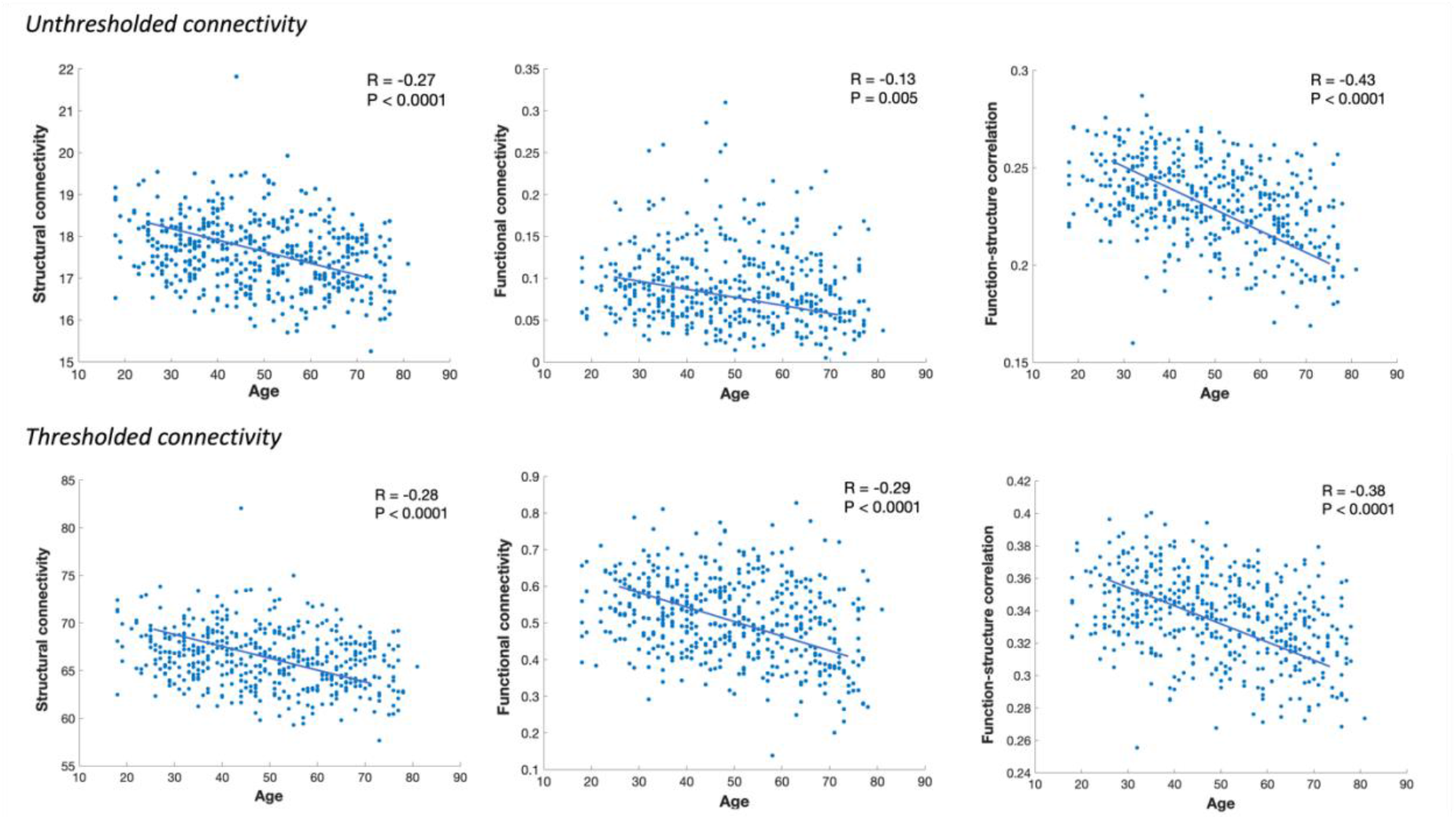
Correlation with age of structural connectivity, functional connectivity and function-structure connectivity correspondence. Each dot represents a participant. Connectivity represents the average connectivity across all connections of that participant. The top row shows the correlation plots of unthresholded structural connectivity and functional connectivity (30135 connections), and the bottom row shows the correlation plots of structural connectivity and functional connectivity thresholded according to structural connectivity (i.e., connections that were 0 in the number of streamlines for ≥50% of subjects were excluded in the analysis, 7934 remaining connections).

The subject-average connection strength of *functional connectivity observed, function-structure connectivity convergence* and *function-structure connectivity divergence* arranged in 8 networks (cortical and subcortical) are shown in **Figure 3**. The subject-average connection strength of *function-structure connectivity convergence* and *function-structure connectivity divergence* when applying a higher structural connectivity threshold (**Supplementary Figure 1**) showed consistency with the main results across networks. The connection patterns of *function-structure connectivity convergence* resembled those of *functional connectivity observed*. The connection patterns of *function-structure connectivity divergence* were generally similar to the *function-structure connectivity convergence* but showed a weaker effect. Connections of the subcortical network itself and with other networks were most significantly weakened in *function-structure connectivity divergence* compared to *function-structure connectivity convergence*. Significantly weakened connections in *function-structure connectivity divergence* were also observed for the visual network.

**Figure 3.**
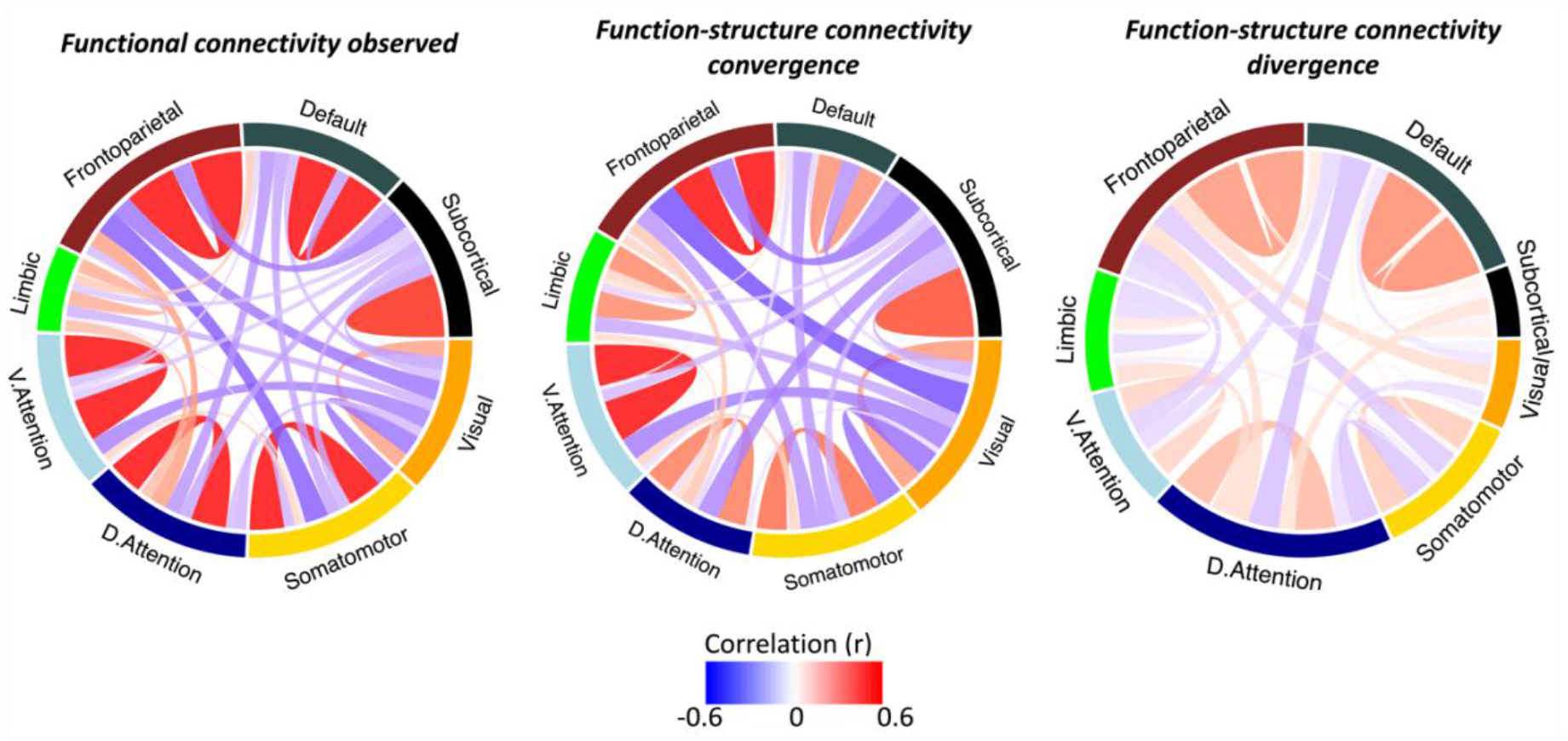
The within- and between-network connection strength of the *functional connectivity observed, function-structure connectivity convergence* and *function-structure connectivity divergence*. Each connection strength link represents the average connectivity across all subjects. Abbreviations: D.attention, dorsal attention; V.attention, ventral attention.

### 3.2 Age effects on connectivity

The effects of *function-structure connectivity convergence* and *function-structure connectivity divergence* on age are shown in **Figure 4**. Overall, *function-structure connectivity convergence* showed stronger effects with age than *function-structure connectivity divergence* in higher-order connections and the subcortical-cortical connections. For *function-structure connectivity convergence*, as age increases, the within-network connectivity generally decreased and between-network connectivity increased, with the exception of limbic-to-transmodal network connections showing a decrease with age. The strongest decrease in within-network connectivity with age was observed within the subcortical network, frontoparietal network, limbic network and ventral attention network. A significant increase in connectivity with age was found in subcortical-to-high-order network connections (e.g., default mode network and frontoparietal network), as well as unimodal-to-transmodal network connections.

**Figure 4.**
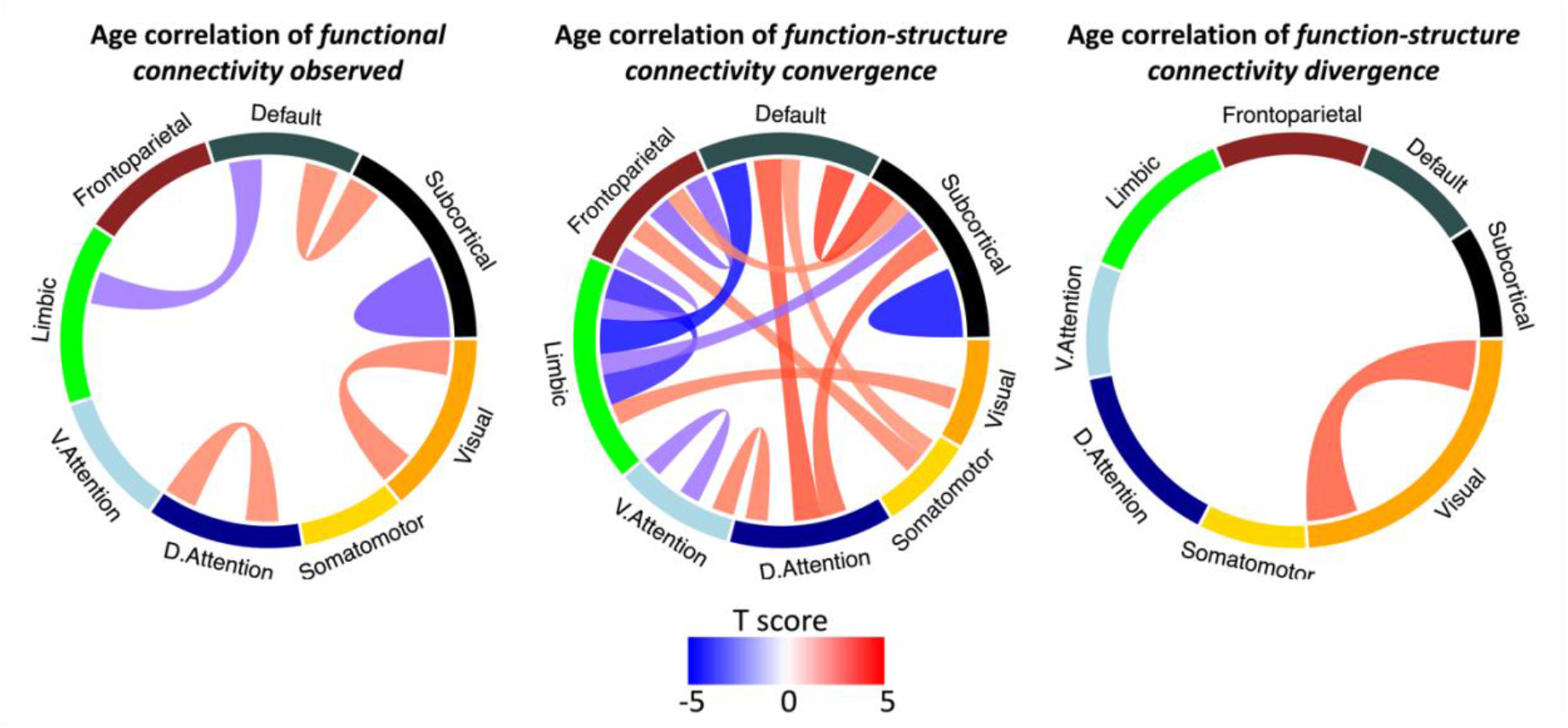
Significant correlations with age of the within- and between-network connections in *functional connectivity observed, function-structure connectivity convergence* and *function-structure connectivity divergence*. Each connection strength link represents the T score of the connectivity variable in the linear regression models: Age ∼ *Functional connectivity observed* + Gender + Head motion; Age ∼ *Function-structure connectivity convergence* + Gender + Head motion; and Age ∼ *Function-structure connectivity divergence* + Gender + Head motion. Only significant connections are shown (FDR corrected *P* < 0.05). Abbreviations: D.attention, dorsal attention; V.attention, ventral attention.

### 3.3 Relationship between connectivity and cognition

In the PLS with cognition, the first component was significant for *functional connectivity observed* (R = 0.24, *P* < 0.0001), *function-structure connectivity convergence* (R = 0.31, *P* < 0.0001) and *function-structure connectivity divergence* (R = 0.19, *P* = 0.002). This component represented overall cognitive performance covering all domains investigated (**Figure 5**). In the multiple linear regression predicting cognitive performance (**Table 2**), we found a significant interaction effect of Age\**Functional connectivity observed* (β = 0.10, *P* = 0.022); a separate model found a similar interaction effect of Age\**Function-structure connectivity convergence* on cognitive performance (β = 0.11, *P* = 0.0079). We observed no such interaction of Age\**Function-structure connectivity divergence* on cognitive performance (β = 0.0074, *P* = 0.86). To help visualize this Age x Network connectivity interaction, scatter plots of **Figure 5** show the correlation between network connectivity and cognition of component 1 from PLS, separated into 3 groups based on subject age (Young: 18-40 years old, *n* = 141; Middle: 41-57 years old, *n* = 142; Old: 57-81 years old, *n* = 142). In older subjects, a positive association with cognition was found in *function-structure connectivity convergence* but not in *function-structure connectivity divergence*. Such increased association with age was found to be significant across all networks for *function-structure connectivity convergence*, with the strongest effects observed between the subcortical and the default mode network, between the subcortical and the frontoparietal network, between the subcortical and the dorsal attention network, between the visual network and the limbic network, between the default mode network and the limbic network, within the limbic network, within the ventral attention network and within the subcortical network.

**Table 2.** Results of multiple regression predicting cognitive performance.

| **Model: Cognition ~ Network connectivity\*Age + Gender + Head motion** |  |  |  |  |  |  |  |  |  |  |
| --- | --- | --- | --- | --- | --- | --- | --- | --- | --- | --- |
| Predicting variables | Network connectivity |  | Age |  | Network connectivity*Age |  | Gender |  | Head motion |  |
| | $\beta$ | $P$ | $\beta$ | $P$ | $\beta$ | $P$ | $\beta$ | $P$ | $\beta$ | $P$ |
| ***Functional connectivity observed*** | 0.093 | 0.029 | -0.44 | < 0.0001 | 0.10 | 0.022 | 0.039 | 0.33 | -0.12 | 0.0058 |
| ***Function-structure connectivity convergence*** | 0.085 | 0.051 | -0.47 | < 0.0001 | 0.11 | 0.0079 | 0.040 | 0.32 | -0.11 | 0.012 |
| ***Function-structure connectivity divergence*** | 0.086 | 0.043 | -0.45 | < 0.0001 | 0.0074 | 0.86 | 0.048 | 0.25 | -0.12 | 0.010 |

**Figure 5.**
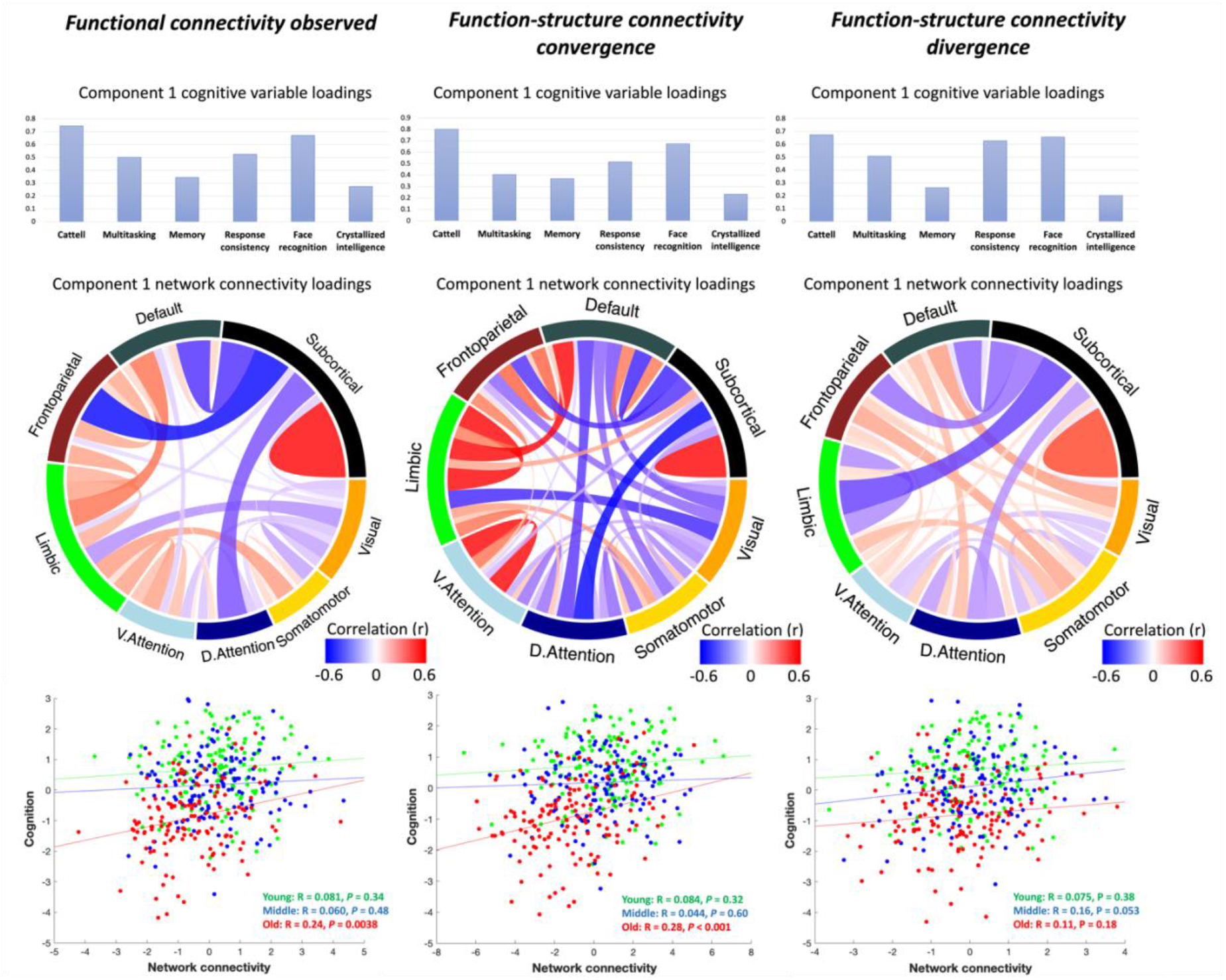
Comparison of the significant components in PLS between cognitive variables and *functional connectivity observed, function-structure connectivity convergence* and *function-structure connectivity divergence*. The top row shows the loadings of cognitive variables, the second row shows the connectivity loadings between and within networks, and the bottom row shows the scatter plots of correlation between network connectivity (i.e., connectivity scores) and cognition (i.e., cognition scores). Each dot represents a participant in the scatter plots. The age effect was analyzed as a continuous variable but for ease of visualization it is illustrated by separation into 3 age groups (Young: 18-40 years old, *n* = 141; Middle: 41-57 years old, *n* = 142; Old: 57-81 years old, *n* = 142). In older subjects, an increased association with cognition was found in *function-structure connectivity convergence* but not in *function-structure connectivity divergence*. Abbreviations: D.attention, dorsal attention; V.attention, ventral attention.

## 4. Discussion

There are two principal results of this study. First, we confirmed that maintaining successful performance in cognitive tasks across the adult lifespan is increasingly dependent on the functional integration across interconnected networks. Second, we tested the brain maintenance theory (Cabeza et al., 2018), distinguishing cognitively-relevant variability in functional connectivity across the lifespan that is shared with structural connectivity (i.e., *function-structure connectivity convergence*), from variability that is independent of structural connectivity (i.e., *function-structure connectivity divergence*). *Function-structure connectivity convergence* showed stronger associations with age and cognition than *function-structure connectivity divergence*. As people get older, cognition across multiple domains depends on functional connections that correspond to the structural connections, especially those involving high-order cortical networks and subcortical networks. The results suggest that maintaining cognitive function in old age is at least partially attributable to the maintenance of functional network connectivity facilitated by structural connectivity (i.e., brain maintenance), with little evidence for the contribution from functional connectivity that is independent of age-related loss in structural connectivity.

There is extensive evidence that functional and structural connectivity are closely related to each other (Deco, Jirsa, & McIntosh, 2011; Hermundstad et al., 2013; Vazquez-Rodriguez et al., 2019). The correspondence between functional and structural connectivity may be due to structural connectivity patterns constraining functional communications between pairs of directly connected regions (Deco et al., 2011; Honey et al., 2009). However, this relationship may change with age and disease. Previous studies have shown that maintaining functional connectivity profile is increasingly relevant for older adults to maintain cognitive functions across different cognitive domains (Bethlehem et al., 2020; Bruffaerts et al., 2019; Guardia et al., 2022; Tibon et al., 2021; Tomassini et al., 2022; Tsvetanov et al., 2016; Tsvetanov et al., 2018). Given the age- and cognition-related correspondence between functional connectivity and structural connectivity, we hypothesized that the increased reliance on functional connectivity for cognition in ageing is dependent on structural connectivity. We first demonstrated that *function-structure connectivity convergence* showed an age-related pattern similar to the pattern of resting-state functional connectivity established in previous literature: within network connections decrease and between network connections increase as age increases (Bethlehem et al., 2020; Betzel et al., 2014; L. Geerligs et al., 2014; Guardia et al., 2022). Overall *function-structure connectivity convergence* showed stronger correlations with multiple cognitive domains in comparison with *functional connectivity observed* and *function-structure connectivity divergence*. We then provided evidence supporting the increased reliance on the *function-structure connectivity convergence* for cognition in older subjects.

Cognitive functions including fluid intelligence, multitasking, memory, response consistency, face recognition and crystallized intelligence, showed age-dependent correlation with *function-structure connectivity convergence* but not *function-structure connectivity divergence*. In particular, cognitive performance was positively correlated with *function-structure connectivity convergence* in older subjects only, not in young and middle age subjects. Such increased reliance on the similarity of structural and functional networks with age was found to be significant across all networks, with the strongest effects observed across high-order cortical networks and subcortical regions. Using multivariate analyses, we have previously demonstrated that the cognitive functions tested in this study were correlated with age and cognition (Guardia et al., 2022; Tibon et al., 2021) and moreover increasingly associated with resting-state effective connectivity profile in older subjects (Bethlehem et al., 2020; L. Geerligs et al., 2017; Tsvetanov et al., 2016). The findings of the present study suggest that the functional network connectivity important for maintaining cognitive performance, especially when people get older, is facilitated by maintaining structural network integrity. The results are consistent with the brain maintenance hypothesis whereby successful ageing is underpinned by maintaining youth-like neural structure and function (Duzel, Schutze, Yonelinas, & Heinze, 2011; Nyberg, Lovden, Riklund, Lindenberger, & Backman, 2012).

It has been proposed that functional reorganization underlies the maintenance of normal cognitive performance in old age, and early stages of diverse neurodegenerative diseases (Cabeza et al., 2018; Dillen et al., 2016; Gregory, Long, Tabrizi, & Rees, 2017; Jones et al., 2016; Simioni, Dagher, & Fellows, 2016; Tsvetanov et al., 2016). An important feature of functional reorganization is the recruitment of brain regions or networks in old adults, despite structural atrophy, leading to better cognitive or behavioural performance (Cabeza et al., 2018). Hence, one might expect cognition to be more dependent on *function-structure connectivity divergence* as age increases if functional reorganization has occurred by increasing the functional connectivity independent of structural changes. However, in this study we found the relationship between cognition and *function-structure connectivity divergence* did not vary with age. This age invariant function-cognition relationship may be more consistent with the brain maintenance theory. A possible explanation for the absence of age-variant relationship between cognition and *function-structure connectivity divergence* is that reorganization might be evident in task-related network connectivity when cognitive demands are more significantly increased, instead of resting-state network connectivity. Second, reorganization might be significant at the prodromal stage of cognitive impairment or neurodegenerative disease (Gregory et al., 2017) but not in healthy ageing subjects with good cognitive reserve by factors such as education; in the present sample, approximately 65% of subjects have at least a degree-level education. We might be able to better test for functional reorganization by performing the same type of analysis (i) at the prodromal stage of dementia or in asymptomatic subjects with a high risk of dementia and (ii) using task-based modulation for functional connectivity.

We found that the connections between subcortical and other cortical networks were weak in *function-structure connectivity divergence*. One interpretation for the differences in patterns of *function-structure connectivity convergence* and patterns of *function-structure connectivity divergence* could be a greater reliance on the strong structural connectivity between subcortical and other regions with increasing age (Cunningham, Tomasi, & Volkow, 2017). In contrast, *function-structure connectivity convergence* showed age decline within the subcortical network and age increase between networks (subcortical-default mode, subcortical-frontoparietal, and subcortical-dorsal attention). *Function-structure connectivity convergence* also showed an age-dependent correlation with cognitive functions between the cortical and subcortical networks. Evidence from previous studies has demonstrated the relevance of subcortical regions, including the thalamus and basal ganglia, with multiple cognitive functions including language, working memory, and numerical cognition (Deldar, Gevers-Montoro, Khatibi, & Ghazi-Saidi, 2021; Moeller et al., 2015). Our study results emphasize the significance of subcortical regions in relation to cognitive functions in normal ageing. While a majority of connectivity studies have focused on cortical regions, it is also important to further understand the connectivity within subcortical regions and their connectivity with other cortical regions, which might be particularly relevant to the maintenance of cognition in ageing.

There are limitations to this study. First, functional connectivity is susceptible to motion and physiological artefacts, such as the non-neural signals due to head motion, breathing and heart rate variability during rs-fMRI scans (Birn, Diamond, Smith, & Bandettini, 2006; Chang et al., 2013; Power, Barnes, Snyder, Schlaggar, & Petersen, 2012). While this is a potential problem for all functional connectivity analyses, we tried to minimize the effect by further processing rs-fMRI data using a whole-brain ICA of single-subject time series denoising method, complemented with a general linear model detrending of the fMRI signal (L. Geerligs et al., 2017). The head motion of each subject was also included as a covariate in all multiple regression models. Secondly, the current analysis focused on mapping structural connectivity patterns based on streamline counts, but multiple empirical and theoretical accounts have identified a relatively weak correspondence between structural and functional matrices (Suarez, Markello, Betzel, & Misic, 2020). Graph theoretical techniques which investigate the use of more derived measures of structural connectivity (e.g., communicability) may offer another solution to help close this gap (Seguin, Mansour, Sporns, Zalesky, & Calamante, 2022). Thirdly, this study was cross-sectional even though the cohort covers the adult lifespan. To accurately understand the effect of ageing, as a longitudinal process, future studies would be strengthened by longitudinal analysis of the *function-structure connectivity convergence* and *divergence* in relation to changing cognition. Lastly, participants in the Cam-CAN cohort were recruited within the Cambridge City (UK) area. Although representative of the local population, further work would be required to study the effects of diversity in socioeconomic, cultural, racial or ethnicity factors.

## 5. Conclusion

In conclusion, we demonstrate that functional connectivity shaped by structural connectivity and functional connectivity independent of structural connectivity show distinct network mappings in relation to age and cognition. Maintenance of cognitive functions in older people depends on functional connectivity supported by stronger structural connectivity, especially those between the high-order cortical networks and the subcortical regions. These results highlight the potential for network- and connectivity-guided targets for maintaining cognition at an older age.

## Data and code availability statement

Data of the Cambridge Centre for Ageing and Neuroscience (Cam-CAN) study are available via the Cam-CAN’s portal (https://camcan-archive.mrc-cbu.cam.ac.uk/dataaccess). The code supporting this study’s findings is available on GitHub: https://github.com/xulinliuxl. Data restrictions may apply in order to preserve participant confidentiality.

## Acknowledgments

The Cam-CAN research was supported by the Biotechnology and Biological Sciences Research Council (grant number BB/H008217/1). This work is supported by the Guarantors of Brain (G101149), the Wellcome Trust (103838), the Medical Research Council (SUAG/092 116768; and SUAG/010 RG91365), European Union’s Horizon 2020 (732592) and the NIHR Cambridge Biomedical Research Centre (BRC-1215-20014, NIHR203312). X.L. is supported by the Cambridge Commonwealth, European and International Trust and the China Scholarship Council. For the purpose of open access, the author has applied a CC BY public copyright licence to any Author Accepted Manuscript version arising from this submission. The views expressed are those of the authors and not necessarily those of the NHS, the NIHR or the Department of Health and Social Care.

## Ethics approval statement

Ethical approval of the Cam-CAN study was obtained from the Cambridge 2 Research Ethics Committee.

## Patient consent statement

Written informed consent was given by all participants.

## Credit author contribution statement

Xulin Liu: study design, analysis, interpretation, writing primary draft. Lorraine K. Tyler: funding, study design, data collection. Cam-CAN: data collection and verification. Simon Davis: analysis. James B. Rowe: funding, study design, interpretation. Kamen A. Tsvetanov: study design, analysis, interpretation. All authors read and approved the final manuscript.

