## Supplementary Figure 1 for "Cognition’s dependence on functional network integrity with age is conditional on structural network integrity"

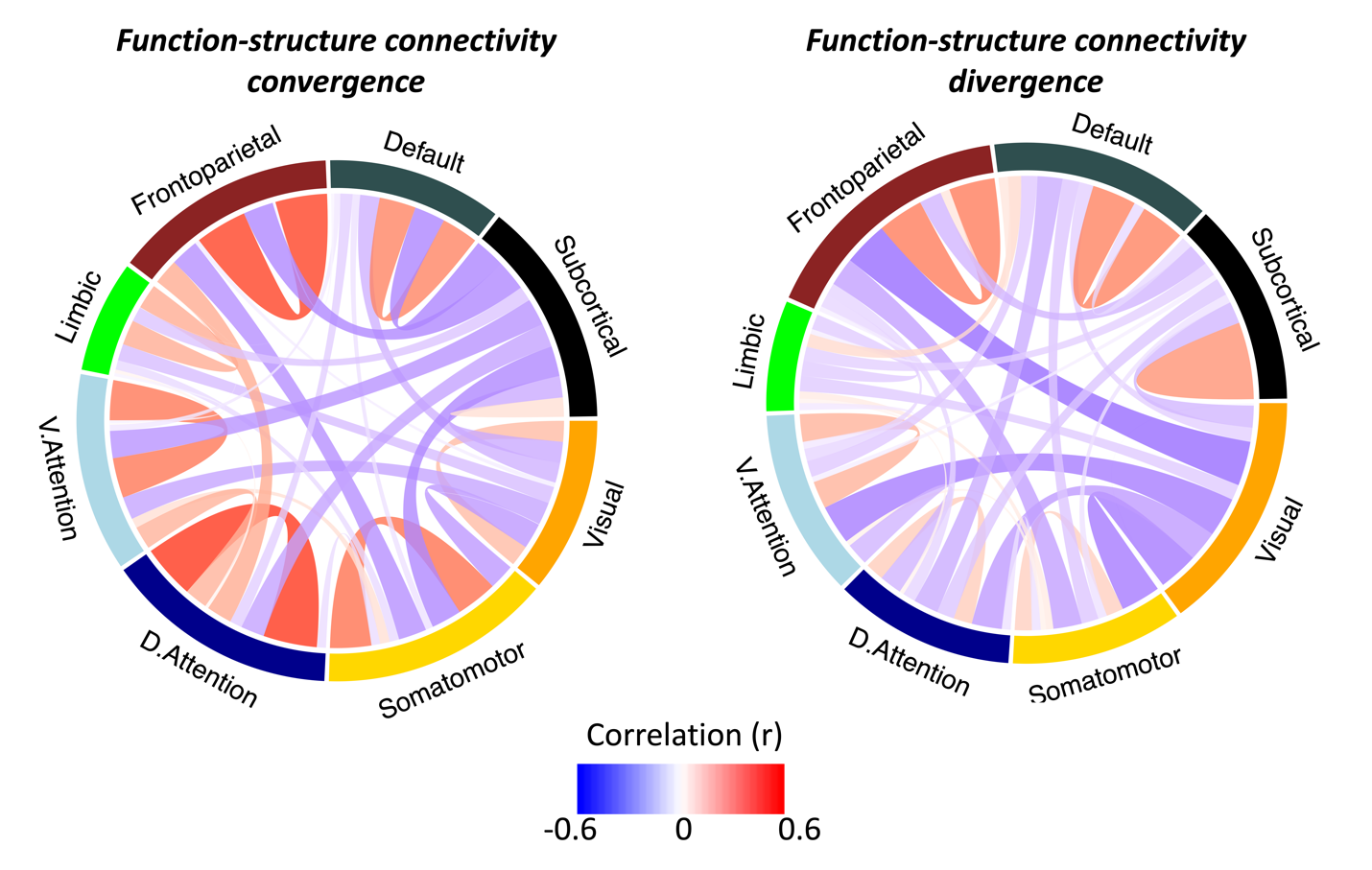

**Supplementary Figure 1**. The within- and between-network connection strength of the functional connectivity which correlated with structural connectivity (*function-structure connectivity convergence*) and the functional connectivity which did not correlate with structural connectivity (*function-structure connectivity divergence*). Connectivity was obtained by using a 90% threshold (i.e., connections that were 0 in the number of streamlines for ≥10% of subjects were excluded in the analysis, 3844 remaining connections). Each connection strength link represents the average connectivity across all subjects. Abbreviations: D.attention, dorsal attention; V.attention, ventral attention.
